# Mesenchymal expression of activated K-ras yields Noonan Syndrome-like bone defects that are rescued by mid-gestational MEK inhibition

**DOI:** 10.1101/634840

**Authors:** Simona Nedelcu, Tatsuya Kobayashi, Monica Stanciu, Henry M. Kronenberg, Jacqueline A. Lees

## Abstract

Activating germline K-ras mutations cause Noonan syndrome (NS), which is characterized by several developmental deficits including cardiac defects, cognitive delays and skeletal abnormalities. NS patients have increased signaling through the MAPK pathway. To model NS skeletal defects and understand the effect of hyperactive K-ras signaling on normal limb development, we generated a mouse model in which activated K-ras^G12D^ was expressed specifically in mesenchymal progenitors of the limb bud. These mice display short, abnormally mineralized long bones that phenocopy those of NS patients. This defect was first apparent at E14.5, and was characterized by a delay in bone collar formation. Coincident mutation of p53 had no effect on the K-ras^G12D^ induced bone defect, arguing that it is does not result from senescence or apoptosis. Instead, our data revealed profound defects in the development of the committed osteoblasts; their appearance is delayed, concordant with the delay in bone collar formation, and they display an aberrant localization outside of the bone shaft. Additionally, we see growth plate defects including a reduction in the hypertrophic chondrocyte layer. Most importantly, we found that *in utero* delivery of a MEK inhibitor between E10.5 and E14.5 is sufficient to completely suppress the ability of activated K-ras to induce NS-like long bone defects in embryogenesis. These data define a critical point in mid-gestation in which elevated MAPK signaling impairs growth plate and bone collar formation and yield NS-like limb defects. Moreover, they offer insight into possible therapeutic strategies for skeletal defects in patients with Noonan Syndrome.

**SIGNIFICANCE STATEMENT:** Noonan syndrome is a genetic condition that is characterized by various developmental defects including skeletal abnormalities that lead to short stature. These patients carry mutations that activate Ras/MAPK signaling. We have generated a mouse model that recapitulates these Noonan Syndrome-like bone defects. Analysis of these animals establishes the developmental window in which bone formation goes awry, and reveals disruption of an early event that is critical for the longitudinal growth of bones. Additionally, we show that treatment with an inhibitor of Ras/MAPK signaling during this key developmental window is sufficient to completely suppress these Noonan Syndrome-like bone defects. This offers possible therapeutic strategies for skeletal defects in patients with Noonan Syndrome.

## INTRODUCTION

Noonan Syndrome (NS) is an autosomal dominant genetic disease that appears with a frequency of 1/1000-1/2500 in newborns (1–3). It is one of a group of developmental disorders, called RASopathies, which result from germline mutations in components of the RAS/MAPK pathway that increase signaling through this pathway (4). Mutations in *PTPN11, SOS1, RAF1* and *KRAS* have all been identified in NS (4). 2% of NS patients have activating *KRAS* mutations (5, 6). *KRAS* encodes a small GTPase protein that functions as a molecular switch to regulate cell proliferation, differentiation and apoptosis by propagating signals from the tyrosine kinase membrane receptors to various intracellular effector cascades including the MAPK pathway.

NS is characterized by defects in various mesenchymal tissues including cardiovascular abnormalities, short stature, craniofacial dysmorphia and skeletal defects (1). Mouse models bearing germline mutations in *PTPN11, SOS1* and *RAF1* have been developed, and these display the spectrum of gross anatomical defects characteristic of NS (7–9). Analysis of these models, as well as an endocardium-specific *PTPN11* mutant mouse strain, has yielded key insight into the underlying basis of the cardiac abnormalities that are the primary cause of death of NS patients (7, 9). Other studies have investigated the nature of the craniofacial defects (8, 10). Most importantly, analysis of these NS mouse models showed that continuous prenatal or postnatal treatment with MEK inhibitors is sufficient to rescue both the cardiac and craniofacial defects (8, 10). Given these findings, MEK inhibitors and other Ras pathway modulators are being evaluated for treatment of NS patients (11).

The skeletal deformities represent a major challenge in the life of NS patients, but their etiology is not well understood. Dual-energy X-ray absorptiometry has shown that bone mineralization is decreased in NS children (12) and analysis of collagen I fragments in urine suggests that this reflects increased bone resorption (13). However, no biopsies or histological sections of NS patients’ bones have been reported. Similarly, the skeletal defects of the *PTPN11, SOS1* and *RAF1* mutant NS mouse models have not been probed. At a broader level, the effect of constitutive MAPK pathway activation on osteoblast and chondrocyte differentiation is an area of active investigation (10, 14). Indeed, the function of Ras/MAPK signaling in osteogenesis is controversial, as some studies suggest it promotes osxteoblast differentiation while others support a suppressive role (14).

Here, we have generated a mouse strain in which activated K-ras^G12D^ is expressed in the mesenchyme of the developing embryo to yield hyperactivation of the Ras/MAPK pathway. These animals display severe skeletal abnormalities that resemble those of NS patients. In particular, these animals have profoundly shorter and wider long bones. These defects are first apparent around embryonic day E14.5, and reflect abnormalities in the growth plate and bone collar. Finally, we show that MEK inhibitor delivery between E10.5 and E14.5 is sufficient to rescue the bone phenotype.

## RESULTS

### Expression of activated K-ras in the mesenchymal lineage causes skeletal defects resembling those of Noonan Syndrome patients

Our goal was to develop a mouse model that phenocopied the mesenchymal defects, and particularly the skeletal defects, of Noonan Syndrome patients by driving hyperactivation of Ras/MAPK signaling specifically within the mesenchymal progenitors of the developing embryo. To achieve this, we used a knock-in mouse strain in which expression of an activating *K-ras* allele, *K-ras*^*G12D*^, is controlled by the CRE recombinase (15). We note that germline *K-ras*^*G12D*^ mutation is not observed in NS patients, presumably because [extrapolating from mouse studies (15)] it results in embryonic lethality. However, we chose *K-ras*^*G12D*^ for this study because it yields potent Ras/MAPK signaling. We crossed the conditional inducible *K-ras*^*G12D*^ mouse with the *Prx1-Cre* transgenic strain (16, 17), which expresses CRE in mesenchymal progenitors of the limb bud from embryonic day 9.5. Using reporter mice, we have previously confirmed that *Prx1-Cre* yields efficient recombination in bone, cartilage, skeletal muscle and fat lineages (18).

At E18.5, we found that *Prx1-Cre;K-ras*^*G12D*^ mutants were present at the expected Mendelian ratio (Table 1). However, these mutants failed to thrive. The vast majority died soon after birth, but rare mutants lived up to, but never beyond, three weeks of age. When compared to their littermate controls, which for this and all subsequent analyses were either *Prx1-Cre* (without *K-ras*^*G12D*^) or *K-ras*^*G12D*^ (without *Prx1-Cre*), the *Prx1-Cre;K-ras*^*G12D*^ mutants were all stunted (Figure 1A). In particular, these animals had profound skeletal abnormalities; the front and hind limbs were greatly shortened, and the fingers and toes were club-shaped (Figure 1A). We note that the hind limbs of *Prx1-Cre;K-ras*^*G12D*^ mutants were paralyzed, impeding their ability to feed from their mothers and likely explaining their failure to thrive. These anatomical defects were observed with complete penetrance. Most importantly, they were highly reminiscent of those occurring in human NS patients (3).

**Table 1.** Viability of *Prx1-Cre;K-ras*^*G12D*^ and *p53*^*c/c*^*;Prx1-Cre;K-ras*^*G12D*^ mice at E18.5.

| Parental genotype | *Prx1-Cre* x *K-ras<sup>G12D</sup>* |  | *Prx1-Cre;p53<sup>c/c</sup>* x *K-ras<sup>G12D</sup>;p53<sup>c/c</sup>* |  |
| --- | --- | --- | --- | --- |
| Offspring genotype | CTL | *Prx1-Cre;K-ras<sup>G12D</sup>* | CTL | *p53<sup>c/c</sup>;Prx1-Cre;K-ras<sup>G12D</sup>* |
| Observed viable | 47 | 17 | 43 | 13 |
| Expected viable | 48 | 16 | 42 | 14 |
| p-value | p> 0.95 |  | p> 0.95 |  |

**Figure 1.**
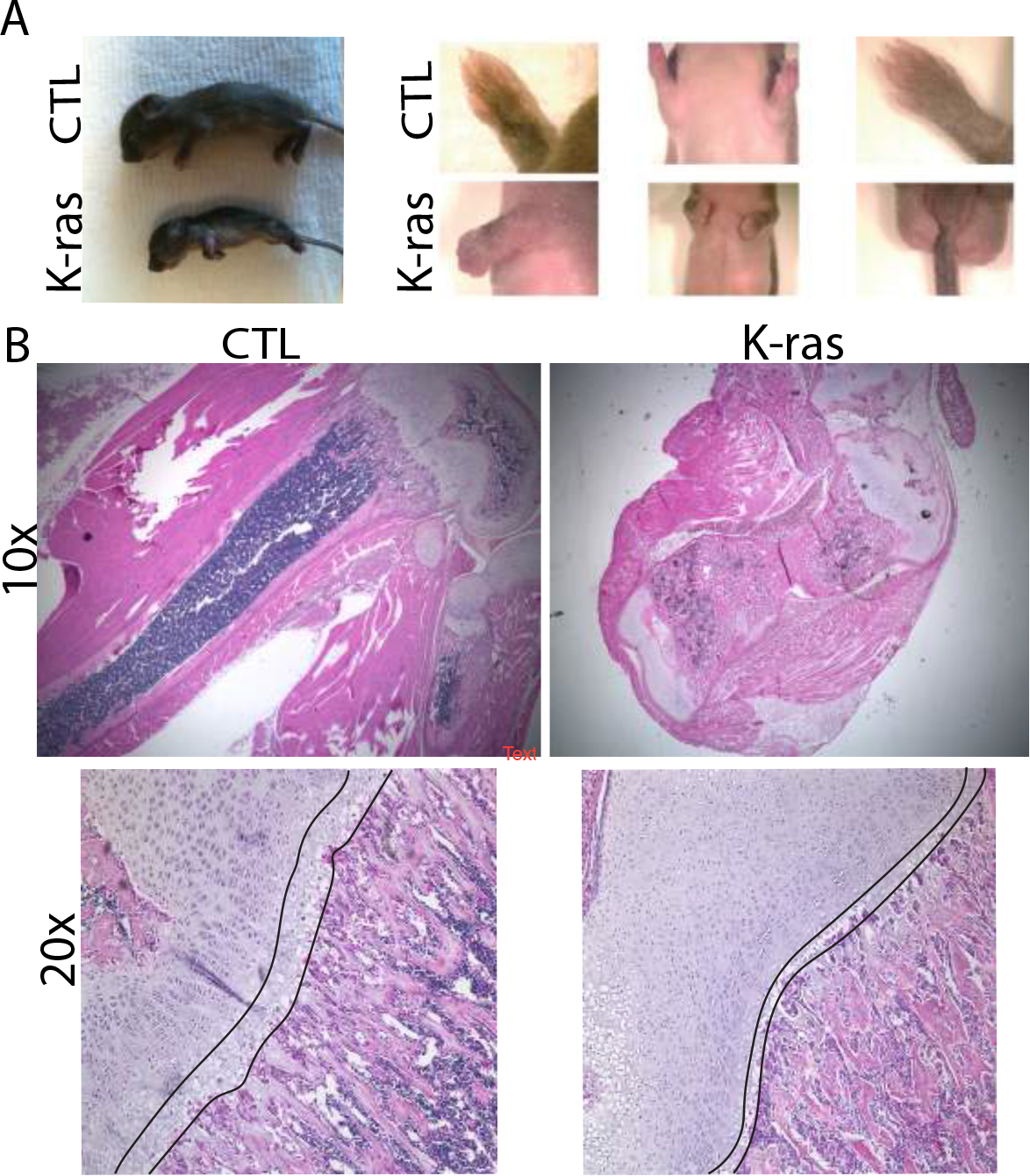
Expression of K-ras^G12D^ in mesenchymal progenitors causes Noonan-like skeletal defects. (A left) Representative 7-days old *Prx1-Cre;K-ras*^*G12D*^ animals are smaller than their wildtype littermates and (A right) their front and hind limbs are greatly shortened and have clubbed feet. (B). H&E staining of representative femur sections from P18 animals shows that the mutant femur is shorter and wider and it has abnormal bone formation at the trabecular region. This, and (B) the higher magnification in the lower panels shows that the mutant growth plate is disorganized and the layer of hypertrophic chondrocytes (indicated by the black lines) is reduced.

We conducted H&E staining of limb sections taken from 18 day old animals, and found that the limbs of *Prx1-Cre;K-ras*^*G12D*^ mutants were significantly shorter (p<0.05, n=10) and wider (p<0.05, n=10) than those of wildtype controls (Figure 1B and data not shown). Additionally, the mutant limbs had abnormal bone tissue spanning the marrow space in the mid-diaphysis and a delay in formation of the secondary ossification center (Figure 1B and data not shown). Indeed, the bone marrow space was almost completely absent in *Prx1-Cre;K-ras*^*G12D*^ mutants. Additionally, cartilage defects were clearly apparent. The growth plates of the long bones were highly disorganized and higher magnification showed that the layer of hypertrophic chondrocytes was much thinner in the mutants than in littermate controls. Notably, these cartilage defects resembled those arising in mice with activating MEK1 mutations (19), suggesting that the bone defects reflect hyperactivation of the MAPK pathway, a conclusion we validate below.

We also conducted a careful analysis of other tissues within *Prx1-Cre;K-ras*^*G12D*^ mutants. We observed no detectable craniofacial abnormalities, even though the mutant heads appeared slightly smaller and *Prx-cre* is known to be expressed in the skull mesenchyme (16). We also observed no cardiac defects, which is consistent with the recent report that cardiomyocyte-specific expression of *K-ras*^*G12D*^ did not yield any defects in heart pathology (20). We confirmed that the conditional *K-ras*^*G12D*^ allele was efficiently recombined in all of the known *Prx-Cre* expressing tissues including the long bones, skull, skeletal muscle and adipose tissue but found that the resulting K-ras^G12D^ expression yielded developmental abnormalities only in the long bones (Figure 1 and data not shown). This striking specificity for bone, but not muscle or fat, defects recapitulates that seen in human NS patients with germline RAS/MAPK pathway mutations (4).

To confirm that the skeletal defect results solely from the expression of K-ras^G12D^ expression in bone and cartilage, we also crossed the conditional *K-ras*^*G12D*^ allele with a *Col2-Cre* transgenic strain. This induces Cre expression to the Col2-positive osteochondral progenitors, which give rise to both chondrocytes and osteoblasts in the developing bone (21). As anticipated, the resulting *Col2-Cre;K-ras*^*G12D*^ animals displayed skeletal defects similar to those of the *Prx1-Cre;K-ras*^*G12D*^ mutants, including shorter and wider long bones with increased trabecular bone bridging the marrow space and aberrant growth plates (Supplemental Figure 1).

### NS-like skeletal abnormalities reflect defects in the appearance and localization of osteoblasts

To establish the etiology of the skeletal defects resulting from K-ras^G12D^ expression, we used timed pregnancies to compare skeletal development in *Prx1-Cre;K-ras*^*G12D*^ mutants with those of littermate controls at various stages of embryonic development. Initially, we examined skeletons of E18.5 embryos using whole mount staining with Alizarin Red, which detects calcium deposits within mineralized bone, and Alcian Blue, which detects the cartilage matrix (Figure 2A). As with the postnatal mutant phenotypes, the *Prx1-Cre;K-ras*^*G12D*^ E18.5 embryos clearly displayed major skeletal abnormalities, of which the most striking were both the front and hind limb defects (Figure 2A and B). Quantification confirmed that the mutant humeri and femurs were both shorter (p<0.05, n=10) and wider (p<0.05, n=10) than those of littermate controls. We also conducted Alizarin Red staining of histological sections from E18.5 limbs (Figure 2C). This clearly showed that the site of mineralization was abnormal and the level of trabecular bone was greatly increased (Figure 2C). The defective nature of long bone differentiation was further illuminated by H&E staining of E18.5 sections, which revealed both the abnormal pattern of mineralization and the presence of atypical growth plates within the greatly shortened and thickened bones (Figure 2D, left panel).

**Figure 2.**
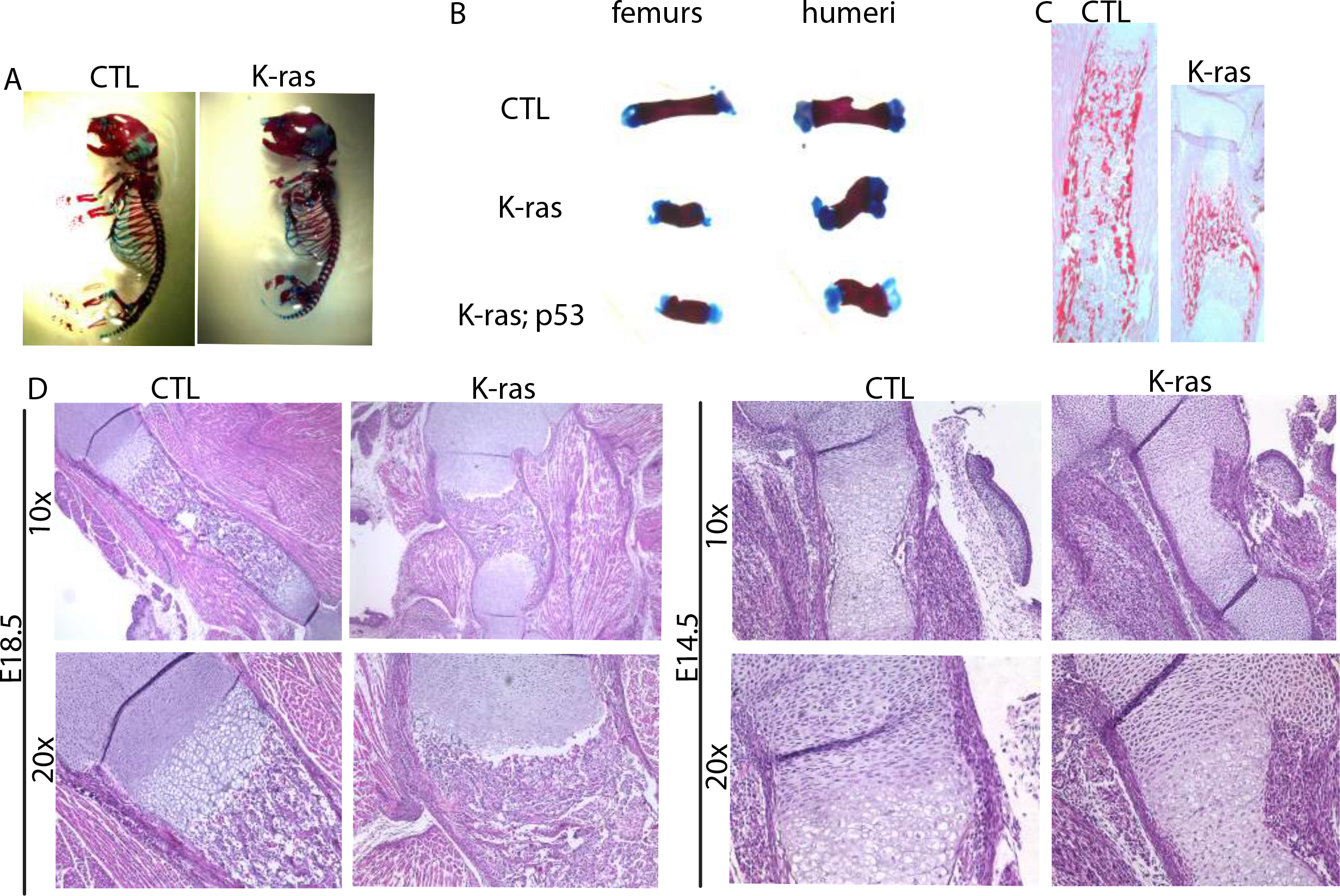
Defects in *Prx1-Cre; K-ras*^*G12D*^ mutants appear early during embryonic development. (A, B) Skeletal staining with Alizarin Red (staining bone) and Alcian Blue (staining cartilage) of E18.5 embryos show that mutants have aberrant skeletal development and that the mutant limbs are much shorter than those of wildtype controls. (C) Alizarin Red staining of representative E18.5 humeri sections shows an increased level of trabecular bone in the mutants. (D) H&E staining of sections from representative E18.5 and E14.5 embryo limbs shows defects at both stages.

To determine when these defects arose, we examined successively earlier development stages. At E16.5, we observed a similar aberrant bone morphology that included both shortened and thickened long bones (Supplemental Figure 2). Analysis of earlier time points traced the earliest morphological indication of bone abnormalities to E14.5 (Figure 2D, right panel). At this stage, the cartilage template of the developing long bones was the appropriate size in *Prx1-Cre;K-ras*^*G12D*^ mutants and the perichondrium also appeared grossly normal. However, while we consistently detected both bone collar formation and the invasion of osteoblasts into the center of the bone in control E14.5 embryos (n = 5/5) both processes were completely absent in the *Prx1-Cre;K-ras*^*G12D*^ E14.5 (n = 5/5) littermates (Figure 2D and data not shown). Since both are present in E16.5 *Prx1-Cre;K-ras*^*G12D*^ embryos, we conclude that hyperactivation of K-ras signaling acts to delay bone differentiation at this critical developmental stage.

It is well established that the activated K-ras can induce senescence and/or apoptosis *in vitro* or *in vivo*, and this is typically p53-dependent. We hypothesized that one, or both, of these activities could account for, or at least contribute to, the stunted bone development in our *Prx1-Cre;K-ras*^*G12D*^ mutants. To test this possibility, we crossed a conditional *p53* mutant allele into our *Prx1-Cre;K-ras*^*G12D*^ model. Remarkably, the deletion of *p53* did not yield any apparent suppression of the *K-ras*^*G12D*^ mutant defects: the *p53*^*c/c*^;*Prx1-Cre;K-ras*^*G12D*^ and *Prx1-Cre;K-ras*^*G12D*^ animals showed the same frequency of perinatal lethality and analysis of E18.5 embryos showed similar long bone defects (Table 1, Figure 2B). Thus, we concluded that the skeletal defects of the *Prx1-Cre;K-ras*^*G12D*^ mice were unlikely to result from either programmed cell death or senescence.

These findings suggested that the activated K-ras was exerting a more direct effect on the differentiation process. Since bone development is influenced by both osteoblast and chondrocyte populations, we conducted *in situ* hybridization (ISH) for Col2, a marker of osteochondral progenitors and also some adult chondrocytes, and Col1, a marker of committed osteoblasts. We examined similar bone sections at E14.5, when the morphological defects were first detected by H&E analyses, and also at E16.5, when the bone defect was striking. Despite the aberrant bone morphology, Col2 staining was appropriately located in the epiphyses of *Prx1-Cre;K-ras*^*G12D*^ mutant bones at both E16.5 and E14.5 (Figure 3A and B). Consistent with this finding, E17.5 humeri of *Prx1-Cre;K-ras*^*G12D*^ and control littermates showed comparable localization of ColX, a marker of hypertrophic chondrocytes (data not shown). In stark contrast, Col1 staining differed greatly between mutant and control animals. At E14.5, Col1 staining was observed specifically in the bone collar of the bones in the control embryos, as expected for this developmental stage (Figure 3A). In contrast, we observed minimal Col1 expression in the bones of *Prx1-Cre;K-ras*^*G12D*^ littermates. This is entirely consistent with the apparent absence of the bone collar in the E14.5 *Prx1-Cre;K-ras*^*G12D*^ H&E sections (Figure 2D). Notably, Col1 was clearly evident at E16.5 in both the *Prx1-Cre;K-ras*^*G12D*^ and control littermates, but the locations were completely different (Figure 3B). Specifically, Col1 staining was detected within the diaphysis of the control embryos, including both the compact and trabecular bones, but was largely concentrated outside of the developing bone structure in the *Prx1-Cre;K-ras*^*G12D*^ mutants. Taken together, these findings argue that hyperactivation of K-ras^G12D^ in the limb mesenchymal progenitors does not disrupt the formation, or localization, of the Col2-positive osteochondral progenitors or committed chondrocytes. Instead, we conclude that K-ras^G12D^ delays the timing of appearance of the Col1-positive, osteoblastic lineages and the bone collar. These Col1-positive cells eventually appear, but accumulate outside, rather than inside, the bone. This has the potential to explain the observed deposition of bone in the lateral, rather than the longitudinal, axis and the consequent formation of wider but shorter bones.

**Figure 3.**
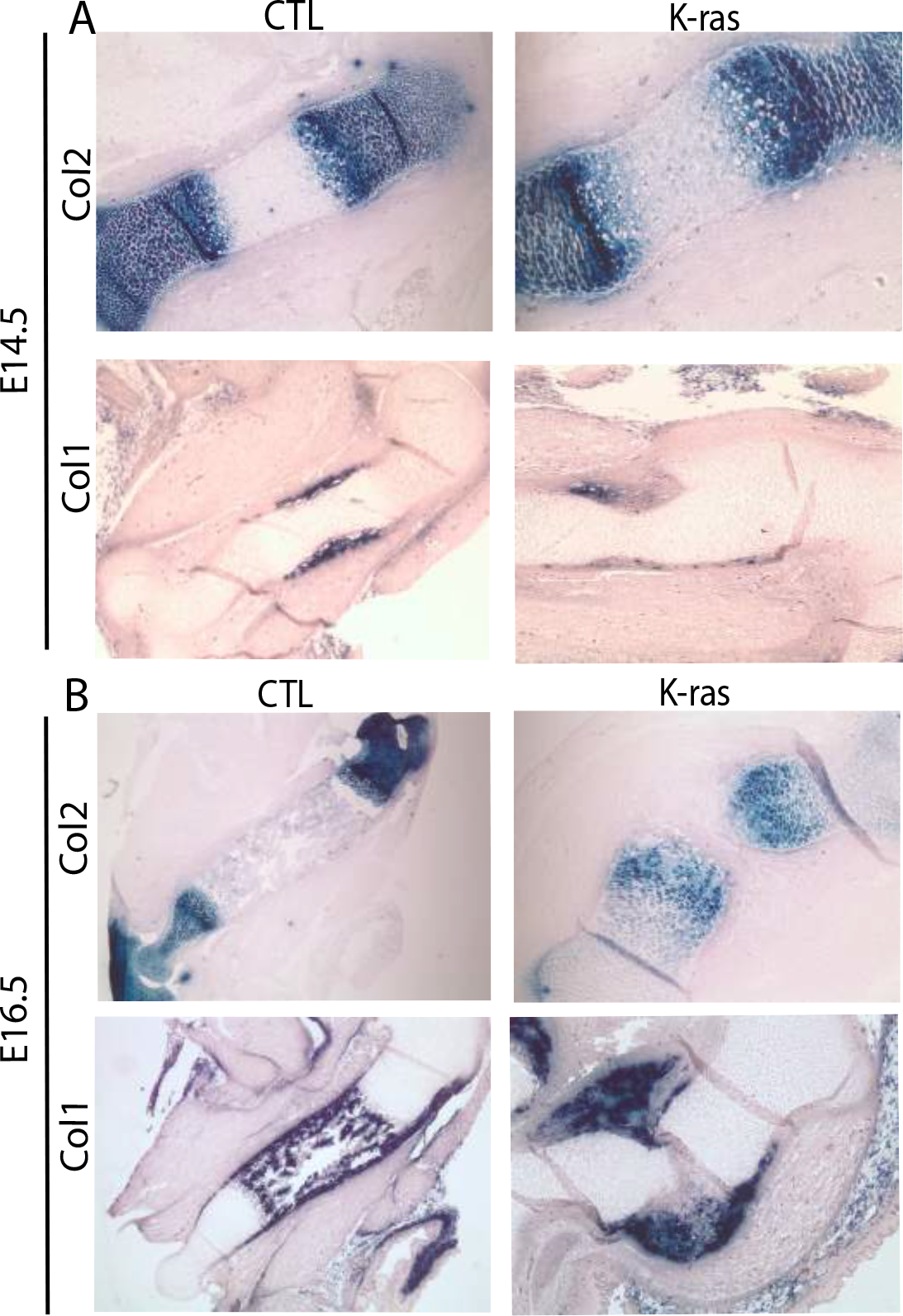
In situ hybridization for bone and cartilage markers on E16.5 and E14.5 limb sections. Representative ISH for Col1 and Col2 in (A) E14.5 and (B) E16.5 embryos shows a disruption of osteoblast and not chondrocyte appearance and location.

### Mid-gestational administration of MEK inhibitor prevents formation of NS-like bone defects

There is considerable evidence to indicate that the defects in RASopathies, including NS, result from deregulation of MAPK signaling (4). Thus, we screened our K-ras^G12D^ mutant long bones for evidence of MAPK activation by conducting IHC for phospho-MEK (pMEK), the active form of MEK. At E14.5, pMEK expression was absent within the majority of the cells within the control humeri (Figure 4A). In contrast, it was present at high levels within most of the bone cells of *Prx1-Cre;K-ras*^*G12D*^ littermates, including the osteoblasts and also the resting, proliferating and hypertrophic chondrocytes (Figure 4A). Thus, the MAPK pathway is hyperactivated at the timepoint we identified as being critical in the development of the NS-like mutant bone phenotype.

**Figure 4.**
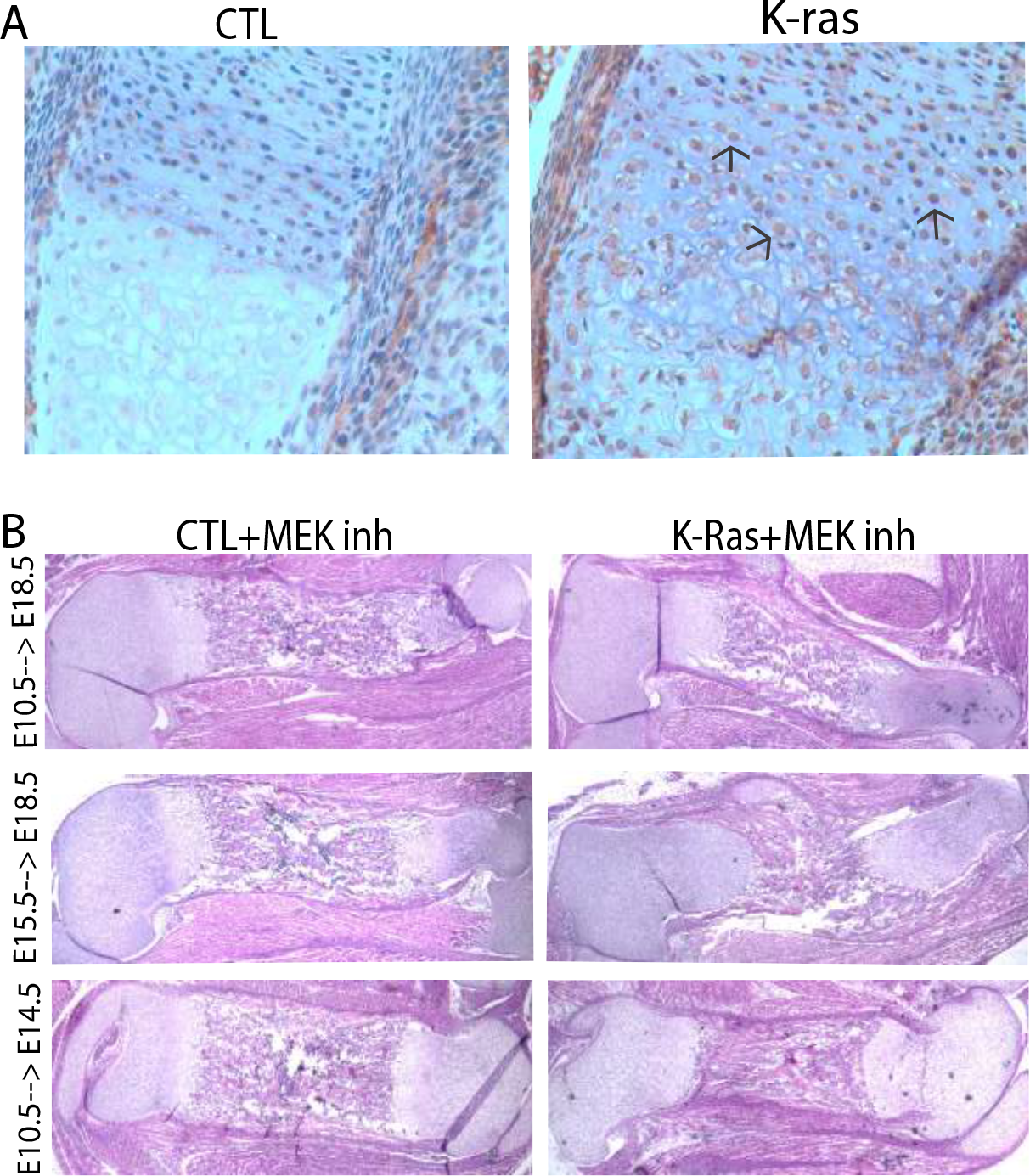
MEK inhibitor treatment during embryonic development rescues the bone phenotypes in *Prx1-Cre;K-ras*^*G12D*^ mutants. (A) IHC for pMEK at E14.5 shows that there is increased pMEK expression in the mutant. Arrows indicate the positive nuclear brown staining. (B) Histological analysis of representative long bones from E18.5 control (CTL) and *Prx1-Cre;K-ras*^*G12D*^ mutant littermate embryos after MEK inhibitor treatment during the indicated time windows.

Given this finding, we wanted to determine whether inhibition of MAPK signaling could suppress this NS-like bone defect. For this, we used the ATP-noncompetitive MEK inhibitor PD0325901 (22). This drug (at 5mg/kg body weight), or vehicle control, was injected into the peritoneal cavity of pregnant mothers for various time periods. Initially, we began treatment at E10.5 and continued daily until E17.5 to encompass the full time period of K-ras^G12D^ activation. The females (n=6 for drug and 4 for vehicle control) were sacrificed at E18.5 and the limbs of the resulting *Prx1-Cre; K-ras*^*G12D*^ mutant embryos (n=4 for drug and 3 for vehicle control) compared to those of littermate controls (n=4 for drug and 3 for vehicle control) by sectioning and H&E analysis. We saw no detectable difference in the long bones of wildtype pups treated with drug versus vehicle control (Figure 4B and data not shown), indicating that MEK inhibition does not modulate normal bone development. In stark contrast, we found that treatment with MEK inhibitor, but never the vehicle control, was sufficient to completely rescue the bone defect of the *Prx1-Cre, K-ras*^*G12D*^ mutants (Figure 4B) and (Supplemental Figure 4). Specifically, bone measurements (Table 2) showed that the PD0325901-treated, mutant limbs were of normal length (p>0.05, n=5) and width (p>0.05, n=5), and their levels of mineralization and of trabecular bone were now comparable to wildtype controls. Thus, we conclude that MAPK signaling is the primary cause of the bone growth defect in the *Prx1-Cre, K-ras*^*G12D*^ mice, and that MEK inhibition is an effective treatment.

**Table 2.** Average femur length and width of CTL and K-ras mice treated with MEK inhibitor for the indicated times (*p<0.05).

| Treatment period | Size (mm) | CTL + MEK Inh | K-Ras + MEK Inh |
| --- | --- | --- | --- |
| E10.5 → E18.5 | Length | 4.23 ± 0.5 | 4.09 ± 0.6 |
|  | Width | 0.7 ± 0.09 | 0.8 ± 0.07 |
| E15.5 → E18.5 | Length* | 4.4 ± 0.7 | 2.1 ± 0.5 |
|  | Width* | 0.6 ± 0.08 | 1 ± 0.06 |
| E10.5 → E14.5 | Length | 4.1 ± 0.3 | 4.2 ± 0.9 |
|  | Width | 0.8 ± 0.07 | 0.7 ± 0.07 |

We wanted to determine whether shorter periods of drug treatment could also suppress the bone defects. Thus, we conducted additional experiments using treatment only between E15.5 and E18.5 or E10.5 and E14.5. Again, embryos were collected at E18.5 and their limbs analyzed. As with the longer treatment, bone development was unaltered in wildtype pups exposed to drug in either time window (Figure 4B and data not shown). For the E15.5 to E18.5 treatment window, the *Prx1-Cre, K-ras*^*G12D*^ limbs displayed the full spectrum of NS-like defects in the case of both vehicle (n=5) and drug (n=3) treated animals, as evidenced by the presence of shorter (p<0.05, n=5) and wider (p<0.05, n=5) limbs (Table 2). In contrast, the mutant bone defect was completely rescued in animals treated from E10.5 to E14.5 with MEK inhibitor (n=5) but not vehicle (n=3) (Figure 4B), (p>0.05, n=5) and width (p>0.05, n=5) (Table 2). Taken together, these experiments define a critical time in bone development, between E10.5 and approximately E14.5, in which upregulation of MAPK pathway signaling is sufficient to derail the process of bone development and yield long bone defects that phenocopy those of NS patients.

## DISCUSSION

In this study, we have generated mouse models in which activation of K-ras^G12D^ in either the mesenchymal progenitors of the limb bud, or the Col2-positive osteochondral progenitors, recapitulates the stunted long bones encountered in patients with Noonan Syndrome. We show that the bones are greatly shortened and thickened, and they display abnormal mineralization in both primary and secondary ossification centers and a stark increase in the density of the trabecular bone. We also see disorganization of the growth plate, and a thinning of hypertrophic chondrocyte layer. The presence of both bone and cartilage abnormalities established the presence of defects in both osteoblast and chondrocyte differentiation.

It is well established that there is crosstalk between these two populations, which is essential for their appropriate differentiation (23). Thus, the activated K-ras in our model system could have a direct effect on one or both cell lineages. Two observations argue for involvement of a chondrocyte defect. First, Murakami and Crombrugge have reported that chondrocyte-specific expression of constitutively active MEK1 yields a cartilage defect that closely resembles the one in our model (19). Second, we did not observe any detectable defects in the differentiation of calvarial osteoblasts, which occurs via a chondrocyte-independent process called intramembraneous ossification. Indian Hedgehog (Ihh) and patched (Ptch) represent the major signaling mechanism between chondrocytes and osteoblasts. Thus, we examined the expression of these markers in the long bones of E14.5 and E16.5 *Prx1-Cre;K-ras*^*G12D*^ mutants and control littermates by in situ hybridization but did not detect any significant differences (data not shown). Although we cannot rule it out, this finding argues against a major defect in in these markers between *Prx1-Cre;K-ras*^*G12D*^ or control littermates. This argues against a defect in the signaling between osteoblast and chondrocyte populations. Given this finding, we favor the idea that activated *K-ras*^*G12D*^ expression is disrupting the behavior and/or timing of differentiation of a shared osteoblast-chondrocyte progenitor.

Our phenotypic analysis tracked the origin of the K-ras^G12D^ mutant bone defects to around E14.5. At this time, the osteoblasts were largely absent, coincident with the absence of the bone collar. Moreover, when the osteoblasts did appear, they accumulated in an aberrant location surrounding, rather than inside, the bone shaft. Interestingly, bone collar osteoblasts are known to originate from Col2-positive osteochondral progenitors, which migrate to the site of the future bone collar and differentiate into Col1-positive osteoblasts (24). Given this, we postulate that the activated K-ras^G12D^ could impair the migration of the Col2-positive osteochondral progenitors, causing them to differentiate in the wrong location. This migration defect hypothesis is supported by prior analyses of zebrafish models for Noonan and Cardio-Facio-Cutaneous syndromes, which showed that expression of mutant RAS, BRAF or MEK disrupts cell movement defects during gastrulation (25, 26). Notably, regardless of the underlying mechanism(s), we believe that the inappropriate positioning of the osteoblasts is responsible for the aberrant deposition of bone in the lateral, rather than the longitudinal, axis.

In addition to elucidating the etiology of NS bone defects, our study provided insight into their potential treatment. Specifically, we found that delivery of the MEK inhibitor PD0325901 *in utero* was sufficient to completely prevent the NS-like long bone defects up to E18.5. Indeed, this treatment was effective when limited to a relatively small time window between E10.5 and around E14.5. This yields two key conclusions. First, it shows that MAPK pathway activation is critical for the development of the NS-like mutant bone phenotype. Second, it proves that the bone developmental defects result from errors that occur during the establishment of bone commitment, presumably the defect in osteoblast recruitment and bone collar formation discussed above.

It is important to note that our data do not establish whether the bone defects are a direct consequence of elevated MAPK signaling, or whether it reflects a compensatory event. Specifically, prior studies have established that MAPK signaling can induce negative feedback loops including the expression of Sprouty family members (27), which are direct inhibitors of RAS and have been shown to modulate bone development (28). Moreover, treatment with MEK inhibitors has been shown to suppress these negative feedback loops in the context of K-ras mutant tumors (29). Additionally, since we examined the PD0325901-treated embryos at E18.5, we cannot preclude the possibility that the presence of activated K-ras^G12D^ signaling post-birth might lead to the formation of other bone defects in neonates/adult animals. We tried to address this issue by allowing treated mothers to give birth. Unfortunately, almost all of the pups died irrespective of their genotype, and irrespective of whether the mother was injected with MEK inhibitor or vehicle control. Thus, we believe that injection scheme itself causes the mothers to reject their pups, and additional experiments will be required to address this question.

Over the past decade, the development and performance of clinical trials in patients with Noonan Syndrome has rapidly increased (11). Many of these trials aim to modify signaling through the aberrantly activated Ras/MAPK pathway. MEK inhibitors have been tested in preclinical mouse models, including PTPN11 and RAF mutant models of Noonan Syndrome and an FGFR model of Alpert Syndrome, and shown to suppress both the cardiac and craniofacial defects (8, 10, 30) Our study now shows that such treatments, if delivered at the time of critical bone development, would also successfully rescue the skeletal defects of NS patients.

## MATERIALS AND METHODS

### Mouse husbandry and drug treatment

Animal procedures followed protocols approved by MIT’s Committee on Animal Care. *K-ras*^*G12D/+*^, *Prx1-Cre* and and *Col2-Cre* animals were maintained on a mixed genetic background. PD0325901 (Sigma-Aldrich) was dissolved in DMSO at 50 mg/ml and diluted to 0.5 mg/ml in vehicle (0.5% hydroxypropyl methylcellulose with 0.2% Tween 80). Mice were injected i.p. with drug (5 mg/kg BW) or vehicle control daily for the indicated times.

### Tissue analyses

For skeletal staining, skin and eviscera were removed from embryos and the remaining tissue placed in 95% ethanol for 4 days, acetone for 3 days, and ethanol containing 0.015% Alcian Blue 8GX (Sigma), 0.005% Alizarin Red S(Sigma) and 5% glacial acetic acid at 37°C for 2 days and then room temperature for 1 day. Tissue was then cleared in 1% potassium hydroxide. For histological analysis, after fixation in PBS with 3.7% formaldehyde, soft tissues were dehydrated via an ethanol series and bone-containing tissue was decalcified in 0.46M EDTA, 2.5% Ammonium Hydroxide pH 7.2. Paraffin embedded sections were cut at 5µm, dewaxed and either stained with H&E or rinsed in water, placed in 2% Alizarin Red S (pH 4.2) for 5 min, dipped 20 times in acetone followed by acetone:xylene *(1:1)*. For pMEK immunohistochemistry, de-waxed slides were incubated in 0.5% H2O2/methanol for 15 min and antigens retrieved using citrate buffer (pH 6.0) in a microwave for 15 min. Slides were blocked in 10% goat serum in PBS for 1 h at room temperature, incubated with anti-pMEK (1:100; Cell Signaling, 9661) in 0.15% Triton/PBS overnight at 4°C, secondary antibodies (1:200; Vector Laboratories) in PBS with 2% goat serum and detected using a DAB substrate kit (Vector Laboratories), and haematoxylin counterstaining. For *In situ* hybridization, tissues were fixed in 4% paraformaldehyde/PBS overnight at 4°C and paraffin embedded sections were cut at 5µm. Non-radioactive ISH, was performed with digoxigenin (DIG)-labeled probes (31, 32) as described previously (32).

## Supporting information

Supplementary Figures-Nedelcu

## ACKNOWLEDGEMENTS

We thank Shigeki Nishimori for the *in situ* hybridization RNA probes for Col1, ColX, Col2, Ihh and Ptch, the Koch Institute Swanson Biotechnology Center for technical support, particularly the Histology and AIPT Cores, and members of the Lees and Kronenberg laboratory for valuable input. This work was supported by NIH/NCI grants to the Koch Institute (P30-CA14051). JAL is a Ludwig Scholar at MIT and SN was supported by a D.H. Koch Graduate Fellowship. HK is supported by NIH DK 56246. TK is supported by NIH AR056645.

