## Supplementary Figures-Nedelcu for "Mesenchymal expression of activated K-ras yields Noonan Syndrome-like bone defects that are rescued by mid-gestational MEK inhibition"

**Supplemental Figure Legends**

**Supplemental Figure 1. *Col2-Cre;K-ras*$^{G12D}$ mice display similar bone defects to *Prx1-Cre;K-ras*$^{G12D}$ mutant animals.** (A) H&E sections and (B) Alizarin Red staining of representative long bones from control versus *Col2-Cre;K-ras*$^{G12D}$ mice at P1.

**Supplemental Figure 2. Long bone defects in E16.5 *Prx1-Cre; K-ras*$^{G12D}$ mutant embryos.** H&E sections of representative humeri of control and *Prx1-Cre;K-ras*$^{G12D}$ mice.

**Supplemental Figure 3. Vehicle injections during embryonic development does not affect the bone phenotypes in *Prx1-Cre;K-ras*$^{G12D}$ mutants.** Histological analysis of representative long bones from E18.5 control (CTL) and *Prx1-Cre;K-ras*$^{G12D}$ mutant littermate embryos after vehicle injections during the indicated time windows indicates no effect on the phenotype.

**Supplementary Figure 1**

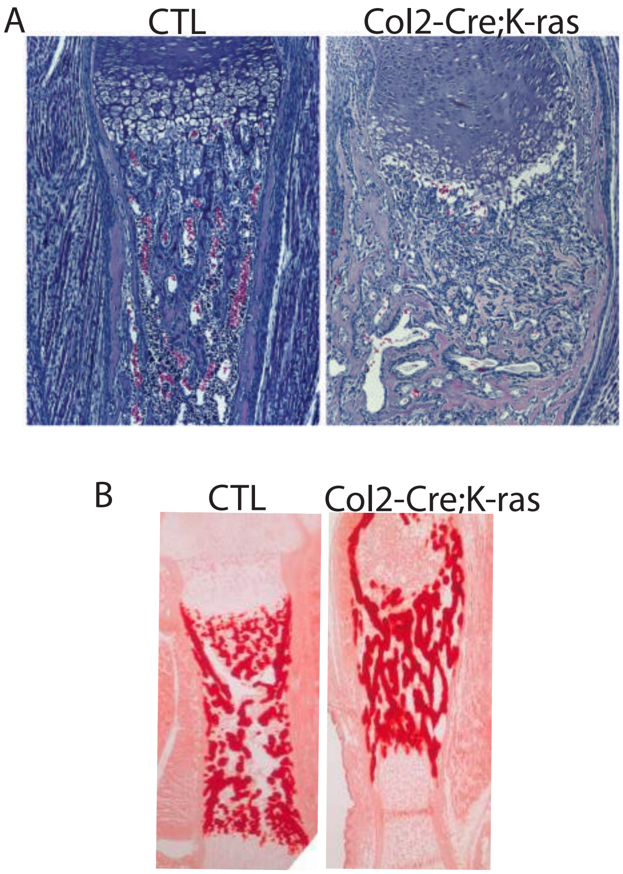

**Supplementary Figure 2**

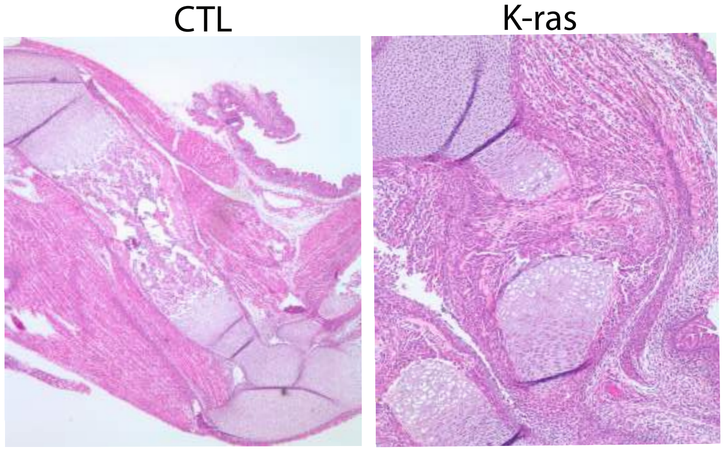

**Supplementary Figure 3**

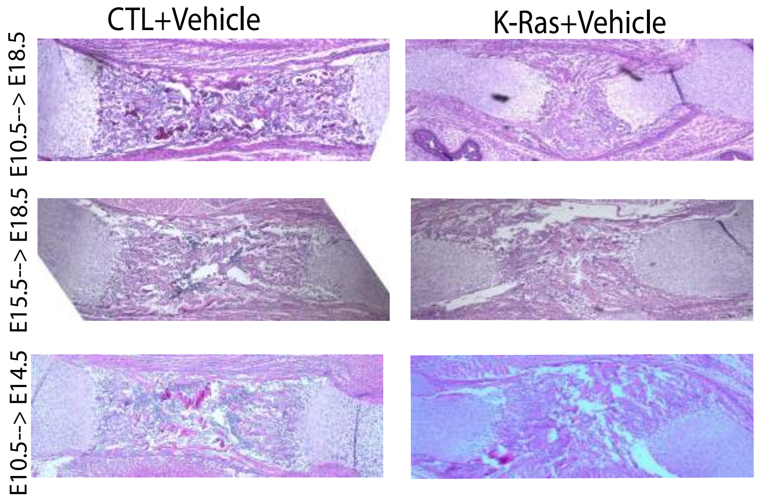
